# Wink or blush? Pupil-linked brain arousal signals both change and uncertainty during assessment of changing environmental regularities

**DOI:** 10.1101/2024.08.07.607112

**Authors:** G. Gesztesi, P. Pajkossy

**Affiliations:** Department of Cognitive Science, Budapest University of Technology and Economics, Budapest, Hungary; Cognitive Medicine Research Group, University of Szeged, Hungary

**Keywords:** uncertainty, change, pupil-linked brain arousal, LC/NA system, entropy, KL divergence

## Abstract

One main cornerstone of adaptive behavior is belief updating, whereby new and unexpected observations lead to the updating of learned associations between events, behaviors and outcomes. This process necessitates the detection of changed environmental contingencies which in turn leads to uncertainty about the environmental regularities. Change and uncertainty are thus inherently linked, and both constructs have been linked to pupil size changes, which might reflect activity in neural networks underlying belief updating. Thus, in our study, we aimed to disentangle the effects of change and uncertainty on pupil-linked brain arousal. We used a probabilistic reversal learning task, where participants had to act according to changing preferences of a fictional character, and used specific cues to independently manipulate the level of change and uncertainty (e.g. the fictional character winked for signalizing change, or his face was blushed to indicate uncertainty). We found that when the cues triggered the same amount of uncertainty, larger levels of change in beliefs led to a transient increase in pupil size during cue processing. In contrast, when the cues signalized a similar amount of change, then increased belief uncertainty was associated with a sustained increase in pupil size, extending in time beyond cue processing. Thus, change and uncertainty exerted independent influence on pupil-linked brain arousal, suggesting the activity of different neural networks, and highlighting the need to disentangle the effects of these overlapping but distinct theoretical constructs.

## 1. Introduction

Several decades of research investigated the psychological and neurobiological associates of non-luminance mediated dilation and constriction of the eye’s pupil (Beatty, 1982; Beatty & Lucero-Wagoner, 2000; Kahneman & Beatty, 1966; Mathôt, 2018). On a psychological level, pupil size changes were associated with mental effort, cognitive control, affective processes and memory (Beatty, 1982; Bradley & Lang, 2015; Goldinger & Papesh, 2012; Kahneman & Beatty, 1966; Oliva & Anikin, 2018; Van der Wel & Steenbergen, 2018; Võ, 2008), whereas on a neurobiological level, noradrenergic, cholinergic and serotonergic influences on pupil size changes were reported (Cazettes et al., 2021; Joshi et al., 2016; Joshi & Gold, 2020; Reimer et al., 2016; Strauch et al., 2022). The most solid evidence has been accumulated for the link between pupil size changes and the firing rate of the Locus Coeruleus (LC), a brain-stem nucleus responsible for the majority of noradrenergic (NA) transmission in the cortex (Aston-Jones & Cohen, 2005; Joshi et al., 2016; Murphy et al., 2014). Although the exact neurobiological background of these links is not yet revealed, pupil size changes can be regarded as an indirect measure of the activity in several brain networks (Joshi & Gold, 2020; Strauch et al., 2022).

Due to this link, pupillometry can be used to imply the neurobiological underpinnings of human decision making: pupil-linked brain arousal was used previously to investigate brain processes underlying uncertainty during perceptual decisions (de Gee et al., 2017; Lempert et al., 2015; Urai et al., 2017) and value-based reinforcement learning (Preuschoff et al., 2011; Van Slooten et al., 2017). Besides, due to its long-established sensitivity to unexpected and surprising events (Lynn, 1966; Sokolov, 1963), the size of the pupil was frequently linked to belief updating (De Berker et al., 2016; Filipowicz et al., 2020; Muller et al., 2019; Nassar et al., 2012; Van Slooten et al., 2017). This term refers to the process whereby beliefs, learned associations between events, behaviors and outcomes, are updated to reflect changes in the environment. Importantly, two related, but separable aspects of such belief updating can be distinguished: On the one hand, the organism has to detect the sudden and unexpected changes in environmental regularities, which cause a mismatch between expected and observed events. On the other hand, the detection of such mismatch leads to uncertainty regarding what are the regularities, which could be used to predict events and outcomes in future. As will be described below, both of these aspects of belief updating have been linked to pupil-linked brain arousal.

### 1.1. Environmental change and pupil-linked brain arousal

The first aspect of belief updating linked to pupil responses can be termed as the detection of *environmental change:* it refers to the process whereby the organism detects a mismatch between expected and observed information, suggesting that the current beliefs about environmental regularities are not tenable and need to be updated. The magnitude of the detected mismatch can be quantified as the probability that the observation signalizes a fundamental change in the environment – this approach was used for example by Yu and Dayan  2005), in their seminal conceptualization of unexpected uncertainty. Alternatively, change related to a surprising event can also be quantified as the information content (i.e. self-information or Shannon surprise) of the event. This is calculated as the negative logarithm of the event probability. This approach can be generalized to the case when the posterior knowledge is also probabilistic. In this case change or surprise can be measured as the Kullback-Leibler divergence (KL divergence, *KLD*) between the prior and posterior distributions. Shannon surprise can be considered as the limiting case of KL divergence, when the posterior probability of the observed event goes to one and the posterior probability of all other options go to zero.

These different assessments of environmental change (i.e. change probability, Shannon surprise and KL divergence) were each found to be correlated with pupil-linked brain arousal and LC/NA system activity (Filipowicz et al., 2020; Nassar et al., 2012; Preuschoff et al., 2011; Zénon, 2019). This suggests that pupil-linked brain arousal is sensitive to the discrepancy between new observations and previous knowledge, and current neurobiological models propose that this link might reflect the involvement of the LC/NA system in resetting or updating neural connections underlying the outdated beliefs (Bouret & Sara, 2004; Sales et al., 2019).

### 1.2. Belief uncertainty and pupil-linked brain arousal

The second aspect of belief updating associated with pupil-linked brain arousal can be referred to as *belief uncertainty* and concerns the level of uncertainty triggered by changing environmental contingencies. If the observed input does not match our current beliefs, then it usually leads to increased uncertainty, because current beliefs cannot be used to reliably describe the environment. The magnitude of such belief uncertainty can be quantified using the entropy (*H*) of the belief probability distribution (see e.g. Bennett et al., 2015; Filipowicz et al., 2020; Kreis et al., 2022; Muller et al., 2019). Importantly, entropy is not only a commonly used measure of uncertainty, but (up to a constant multiplier) it is the only one that satisfies certain reasonable mathematical properties, such as subadditivity in general and additivity for independent events (Aczél et al., 1974).

Similarly to the detection of environmental change, belief uncertainty is also related to pupil-linked brain arousal. Unlike change detection, however, which has been associated with the magnitude of evoked changes, belief uncertainty has been linked to baseline pupil size (Filipowicz et al., 2020; Muller et al., 2019; Nassar et al., 2012; Pajkossy et al., 2023; Van Slooten et al., 2017; Vincent et al., 2019) or the tonic activity of the LC/NA system (Aston-Jones & Cohen, 2005). Furthermore, pupil-linked brain arousal was also found to track the process whereby new observations after change detection enable the formation of new beliefs, leading in turn to a gradual decrease of belief uncertainty: after changes in environmental regularities, baseline pupil size first increases, reflecting the increased uncertainty, and then it tends to gradually decline, once new beliefs are formed and uncertainty decreases (see e.g. Filipowicz et al., 2020; Nassar et al., 2012; Pajkossy et al., 2017, 2018).

### 1.3. Dissociating the effect of environmental change and belief uncertainty in pupil responses: the current study

The interpretation of the above findings linking pupil-linked brain arousal to both environmental change and belief uncertainty is complicated by the fact that change and uncertainty are highly correlated during the process of belief updating: when new observations suggest that currently held beliefs are no longer valid, then the probability of change increases, but at the same time, so does uncertainty regarding the beliefs describing environmental regularities. Thus, it might be challenging to disentangle the effects of environmental change and the resulting belief uncertainty on pupil responses.

Furthermore, to model the probabilistic nature of environmental regularities, several studies used probabilistic stimulus-reward contingencies (e.g. Filipowicz et al., 2020; Nassar et al., 2012; Pajkossy et al., 2023). In such designs, belief updating is a gradual process, whereby accumulation of evidence over multiple trials will determine how change probability and uncertainty unfolds over time, and when and to what extent it will lead to belief updating. The time course of this process, however, is not directly observable, and this complicates the dissociation of these constructs in the pupil responses observed during these trials.

As an additional methodological problem, timing and type of reported pupil responses also differ. Pupil-linked brain arousal is assessed either as stimulus-linked rapid and short dilation of the pupil above its baseline level or as a more slowly evolving change in baseline pupil size (probably reflecting a similar dichotomy in the functioning of the LC - see e.g. (Aston-Jones & Cohen, 2005). As described above, the effects of environmental change were linked to transient phasic pupil dilation triggered by new observations (see e.g. Filipowicz et al., 2020; Nassar et al., 2012; Pajkossy et al., 2023). In contrast, the effect of belief uncertainty was often assessed on a longer time-scale, affecting baseline or tonic pupil size (Filipowicz et al., 2020; Muller et al., 2019; Nassar et al., 2012; Van Slooten et al., 2017). Thus, not only are the above factors co-occurring during adaptation to environmental change, their effect on pupil-linked brain arousal is often measured at separate time points.

To address these theoretical and methodological discrepancies, Pajkossy and associates (2023) aimed to separate the effect of environmental change and belief updating on the same, phasic pupil response. The authors used a probabilistic reversal learning task, where participants were engaged in a guessing game with a fictional character, who hid a stone in either its left or right hand – the task of the participants was to guess the location of the stone repeatedly over multiple trials. The fictional character preferred one of its hands for hiding, so participants had to figure out which of the response options (left or right hand) yields reward more frequently and choose this option repeatedly to maximize reward. To evoke sudden changes in environmental regularities, the identity of the more advantageous response option (i.e. the hand preference of the fictional actor) was sometimes reversed, and then the participants had to adapt their behavior based on the changing pattern of reward frequency. A Bayesian model was used to compute switch probability and belief uncertainty after feedback given to each choice. Switch probability was computed as the probability that the preferred side has been changed from the participant’s last guess, and it was used to assess environmental change (similarly to the concept of unexpected uncertainty proposed by Yu and Dayan, 2005). To quantify belief uncertainty, the entropy of the probability distribution regarding the preferred side was computed. Using multiple linear regression analysis, the authors showed that both factors predicted feedback-evoked pupil dilations, even after controlling for the other’s effects. That is, belief uncertainty was not only associated with baseline pupil size increase (as shown previously by Filipowicz et al., 2020; Nassar et al., 2012), but it also predicted the magnitude of phasic pupil dilation responses: larger pupil responses after stimuli triggered larger uncertainty, even after controlling for the level of change probability associated with this feedback.

A limitation of this study was, however, that the experimental task did not make it possible to independently manipulate the magnitude of change and uncertainty. When using multiple linear regression to separate their effects, only the linear effect of the other variable can be controlled - this linearity assumption, however, might not be true. Moreover, the probability distribution regarding the preferred side during each trial was highly subjective, depending on the participant’s prior expectations. To estimate these distributions, a Bayesian model was used, with prior parameters fitted to the actual choices, but such models can only estimate the participant’s unobservable beliefs with some degree of modeling related noise (see e.g. Wilson & Collins, 2019).

To correct for these limitations, and to extend the results of Pajkossy and associates (2023), in the current study, we modified the design of the study in three important ways. First, changes in environmental contingencies did not occur at random time points, but during specific *test trials*, which occurred only after completing a series of *filler trials* with fixed stimulus-reward contingencies (Figure 1A-B). This design ensured that participants were aware of choice outcome probabilities prior to test trials (i.e. when change in environmental contingencies was introduced). Furthermore, changes in environmental regularities were clearly indicated to the participants, thus they were not to be inferred from feedback over multiple trials. Because of this, we could localize in time when environmental change or belief uncertainty occurred. As a second modification, we used different instructions during test trials (see below) to alter the probability distributions so that the effect of change and uncertainty could be independently investigated. The magnitude of change was assessed by computing the KL divergence value between the reward probability distributions before and after the change, whereas the magnitude of uncertainty triggered by this new information was assessed by the entropy of the posterior reward probability distribution.

**Figure 1.**
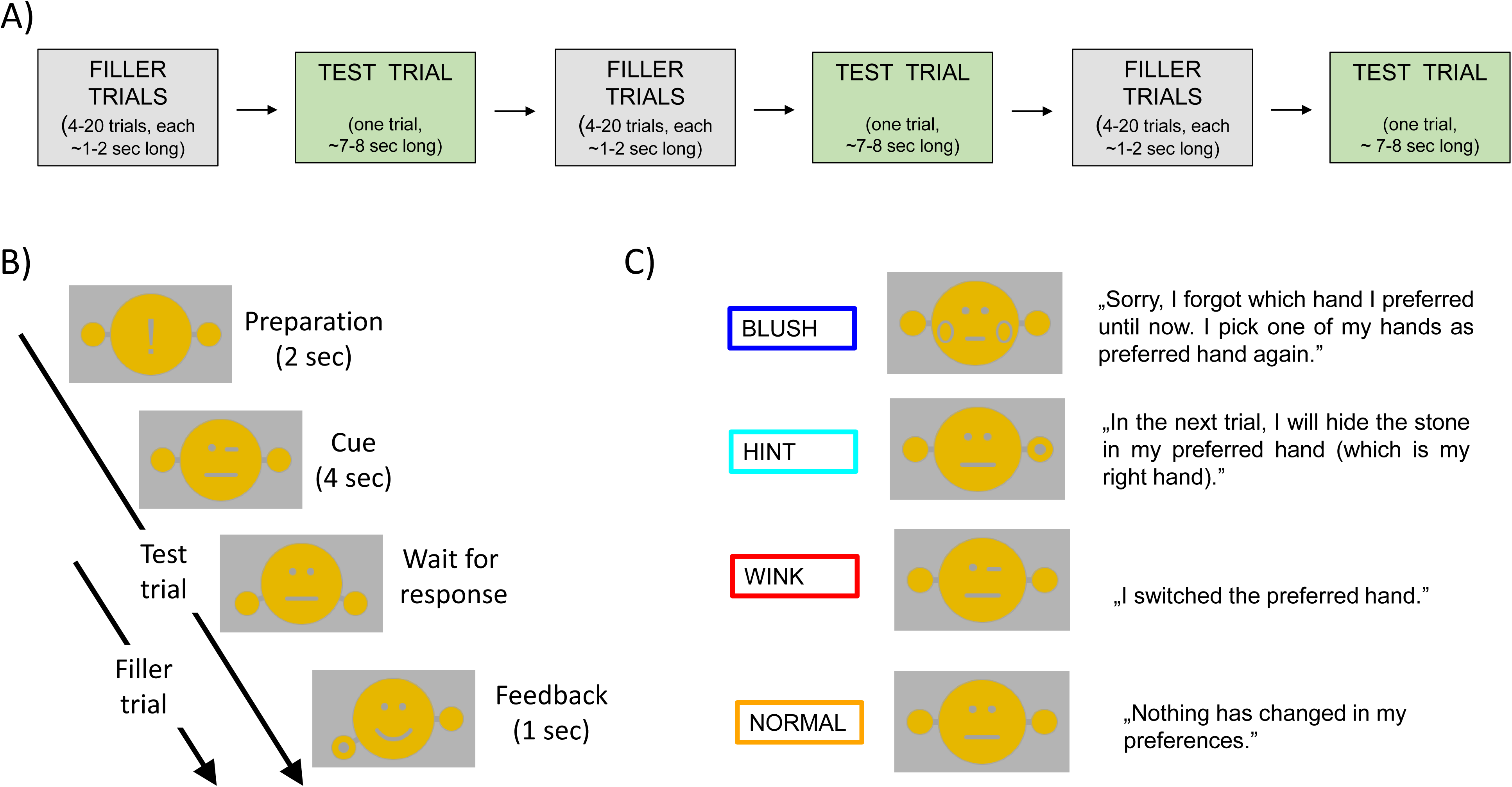
Design of the experiment. A) A series of filler trials and intermixed single test trials were presented to the participant. B) Structure of the trials: during filler trials, the waiting fictional character was presented until the participant gave a response, which triggered a feedback (see bottom two screens). During test trials, first a preparation signal, and then a cue was presented, indicating whether and how the preferences of the fictional actor change (upper two screens). This was followed by the same two screens, which were also presented during filler trials (bottom two screens). C) The cues presented during test trials informing participants whether and how the fictional actor’s preferences change. Screenshots show the visual cue displayed to the participants, the text refers to the meaning conveyed by the specific cues.

Four different types of test trials were used, and the picture of the fictional actor was modified in each trial type, to signalize the nature of the change (see Figure 1C and Table 1). Before the test trials the participants learned the preferred side and knew that it was selected in 80% of the cases (i.e. prior probabilities for the two response options: *p*_1_ = 0.8, *p*_2_ = 0.2). In *blush* trials, the fictional character forgot its preferences, signalized by its face getting blushed, and so the preferred hand was chosen again randomly. Because of this, after this information, participants had no knowledge about which response option was set as the preferred one (i.e. prior probabilities: *p*_1_ = 0.8, *p*_2_ = 0.2; posterior probabilities: *p*_1_′ = 0.5, *p*_2_′ = 0.5, *KLD* = 0.32, *H* = 1). In *hint* trials, the fictional character showed that it will hide the stone in its preferred hand (i.e. prior probabilities: *p*_1_ = 0.8, *p*_2_ = 0.2; posterior probabilities: *p*′_1_ = 1, *p*′_2_, 0, *KLD* = 0.32, *H* = 0). In *wink* trials, the fictional character winked, and this signalized that it changed its preferred hand (i.e. prior probabilities: *p*_1_ = 0.8, *p*_2_ = 0.2; posterior probabilities: *p*′_1_ = 0.2, *p*′_2_ 0.8, *KLD* = 1.2, *H* = 0.72). Finally, in *normal* test trials, participants were signalized that the reward contingencies remained unchanged (i.e. prior probabilities: *p*_1_ = 0.8, *p*_2_ = 0.2; posterior probabilities: *p*_1_′ = 0.8, *p*′_2_ = 0.2, *KLD* = 0, *H* = 0.72).

**Table 1.** Characteristics of the four types of test trials. . Posterior reward probabilities were manipulated so that measures of environmental change and uncertainty enabled the separate investigations of both constructs. KLD: Kullback-Leibler divergence

| Condition | Reward probability |  |  |  | Entropy | KLD | Manipulation |
| --- | --- | --- | --- | --- | --- | --- | --- |
|  | prior - before<br>cue presentation |  | posterior - after<br>cue presentation |  |  |  |  |
| | $p_1$ | $p_2$ | $p_1'$ | $p_2'$ | | | |
| Blush | 0.8 | 0.2 | 0.5 | 0.5 | 1 | 0.32 | Effect of<br>uncertainty -<br>by similar<br>level of<br>surprise |
| Hint | 0.8 | 0.2 | 1 | 0 | 0 | 0.32 |  |
| Wink | 0.8 | 0.2 | 0.2 | 0.8 | 0.72 | 1.2 | Effect of<br>change - by<br>similar level<br>of uncertainty |
| Normal | 0.8 | 0.2 | 0.8 | 0.2 | 0.72 | 0 |  |

Our aim was to test the independent effect of belief uncertainty and environmental change on pupil responses during these test trials. Specifically, the effect of belief uncertainty by similar levels of change was investigated by comparing pupil responses during *blush* and *hint* trials: in these conditions, KL divergence between the prior and posterior distributions measuring change were equal, whereas posterior entropies assessing uncertainty differed. In a similar vein, the effect of environmental change by a fixed level of belief uncertainty was tested by comparing *wink* and *normal* trials: in these conditions, posterior entropy values measuring uncertainty were equal, whereas the KL divergence values assessing change differed (see Table 1).

## 2. Materials and methods

### 2.1. Participants

The participants were graduate and undergraduate students and received monetary compensation for their participation. The size of the initial sample was *N* = 80 (50 females, age range: 18-25, *M_age_* = 21.54, *SD* = 1.69). Due to low pupil data quality, one subject was excluded (see below), thus the final sample size was *N* = 79 (49 females, age range: 18-25, *M_age_* = 21.54, *SD* = 1.70). All subjects had either normal or corrected to normal vision. They were asked to avoid smoking and the consumption of caffeine or alcohol before the experiment on the day of testing. All participants provided informed consent.

### 2.2. Design

#### 2.2.1. Probabilistic reversal learning task

The experimental task was an extended version of a probabilistic reversal learning task. We asked the participants to play a simple game with a fictional character, who repeatedly hid a stone in his left or right hand. The participant then had to make a guess if the stone was hid in the left or the right hand. If this guess was correct, the participant received a point. Otherwise the participant lost a point. The score started from 100. The fictional character always had a preferred hand. He hid the stone with probability 80% in his preferred hand and with probability 20% in his non-preferred hand. These probabilities were known by the participants.

The fictional character appeared on the screen as a small stick figure (see Figure 1). We used two isoluminant colors for its depiction (yellow and dark-gray) on a homogenous light gray background (see Figure 1B-C). The stimulus appeared at the center of the screen and had a small viewing angle of approximately 3 .

An experimental session consisted of 32 blocks. Each block contained a series of up to 20 filler trials followed by a test trial (see Figure 1A). The role of the filler trials was to make sure that the participants learned the preferred hand. During the fillers the preferred hand never changed, and this was also told to the participants. During the test trials, the preferred hand of the fictional character could change depending on the type of the instruction. Both the filler and the test trials were scored. The score was displayed continuously during filler trials and in the feedback phase of the test trials.

A filler trial had two steps: The fictional character showed off his closed hands indicating that the participant could choose. If the participant selected a hand using the left or right arrow button the stick figure opened his hands and the location of the stone became visible. The score was also updated. The feedback was displayed for one second, and then the experiment proceeded with the next trial (see Figure 1B).

The series of filler trials ended if the following two conditions were both true: 1) There were at least 4 more trials in the series when the fictional character hid the stone in his preferred hand as compared to the trials when the stone was in his non-preferred hand. 2) The participant selected the preferred hand consecutively in the last three filler trials. Note that it was not important if the response was correct, only that the participant selected the preferred hand. The first condition ensured that enough information was provided to determine the preferred side with high probability, while the second condition ensured that the participant had indeed learned it. The filler series ended after 20 trials even if the conditions were not met, but in this case the following test trial was omitted from the analysis. The number of trials in a filler series were thus between 4 and 20 (*M* = 7.1, *Mdn* = 6.0, *SD* = 3.6). We omitted 1.4% of the test trials because the above conditions were not met after 20 trials.

The test trials were longer than the fillers to allow capturing the pupillary response (see Figure 1B). They started with a two-second-long preparation phase, during which an exclamation mark was presented, signaling the participant that a test trial was initiated. Then a cue was presented for four seconds indicating the type of the test trial. The remaining selection and feedback steps were identical to the filler trials. After the feedback, the filler trials of the next experimental block started immediately.

There were four different types of test trials: *normal, wink, blush* and *hint*. The meaning of these types was fully explained to the participant before the experimental session. Depending on the type of the test trial sometimes the preferred hand was changed. This happened before hiding the stone, so the rewarded option in the test trial was selected according to the new preferences. The *normal* test trials were similar to the fillers: the preferred hand remained unchanged. In *wink* test trials, the stick figure winked at the beginning of the trial, and this signaled that the preferred and non-preferred hands were switched: the previously non-preferred hand became preferred and the previously preferred hand became non-preferred. In *blush* test trials, the stick figure blushed at the start of the trial, indicating that he ‘forgot’ the preferred hand. In this case the new preferred hand was selected randomly with equal (50%) probabilities. This hand then also became the preferred side for the next block of filler trials. In *hint* test trials the stick figure showed the actual hiding location of the stone at the beginning of the trial, which was the preferred hand. There were eight test trials from each of these four types, resulting in 32 experimental blocks (a set of maximum 20 filler trials followed by a test trial). The order of the different types was randomized, but the same type never occurred three times in a row.

Importantly the cues were constructed to allow independent manipulation of uncertainty and change (see Table 1). Uncertainty was measured as the posterior entropy of the rewarded option, while change was measured by the KL divergence of the posterior reward probability distribution from the prior reward probability distribution. That is if in a test trial the prior (before cue presentation) reward probabilities were *p*_1_ and *p*_2_ and the the posterior (after cue presentation, but before feedback) reward probabilities were *p*′_1_ and *p*′_2_, then uncertainty in that trial was calculated as 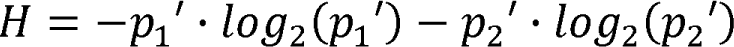 and change in that trial was calculated as 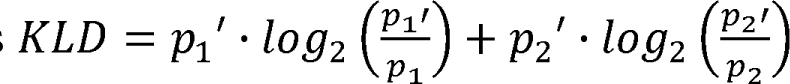. Note that both entropy and KL divergence are assumed to be continuous, so in *hint* trials the term corresponding to the zero posterior probability option is zero. Consequently two of the test trial types, *normal* and *wink* had equal amounts of uncertainty (*H* = 0.72), but different amounts of change (*KLD* = 0 / *KLD* = 1.2 respectively). For the *blush* and *hint* trial types the uncertainty values were different (*H* = 1 / *H* = 0) but the amount of change was the same (*KLD* = 0.32).

#### 2.2.2. Visual Control Task

To make sure that differences in the pupillary responses observed during the test trials are due to differences in belief updating and not due to the low-level visual features of the presented stimuli, each subject participated in a separate visual control task. Importantly this control task was conducted prior to the probabilistic reversal learning task, before any information was given to the participant about the meaning of the stimuli.

The visual control task had 40 trials, each cue type was shown 10 times in randomized order. In each trial first the start trial sign was displayed for 2 seconds, then one of the visual cue stimuli (*wink, normal, blush* or *hint*) for 4 seconds. No task was given to the participant other than looking at the center of the screen where the stimuli were presented. The recorded pupil data were processed and analyzed the same way as described for the experimental sessions.

### 2.3. Procedure

The experiment was conducted in a dimly lit laboratory with constant background illumination. After arriving, participants were seated in front of the computer screen and the eye-tracker. Their head was placed on a chin rest facing a 23-inch monitor with a viewing distance of about 60 cm. First, participants completed the visual control task, which was followed by the probabilistic reversal learning task. Both tasks started with the calibration of the eye-tracker. After finishing both tasks, participants were debriefed.

### 2.4. Pupil Data Acquisition and Processing

The pupil of both eyes were captured using an SMI RED500 eye tracker with 500 Hz sampling rate. The recorded data were preprocessed. Pupil sizes of the left and right eyes were first averaged. Blinks were identified using the data-processing software of SMI, and data points during blinks were removed. Data points more than 3 standard deviations below or above the subject’s mean were deemed outliers. Such outliers and 40 milliseconds before and after the outliers were also deleted.

The remaining data was resampled to 50 Hz, and the missing data points were interpolated linearly. Finally we smoothed the pupil data using a Savitzky-Golay filter. The parameters of the filter were frame size: 9 and polynomial order: 4.

The analysis focused on pupil size changes in the test trials. Trials with more than 25% interpolated data points were excluded from the analysis. Test trials were also excluded if the preceding series of fillers ended after 20 trials, but the participant did not learn the preferred side (see above for the criteria to end a series of filler trials and proceed to the next test trial).

One participant was completely excluded from the analysis (both in the reversal learning and the visual control task) because less than four non-excluded trials remained for at least one experimental condition due to poor quality pupil data. The mean proportion of excluded test trials for the other participants was *M* = 5.1% (*Mdn* = 3.1%, range: 0.0% - 37.5%, *SD* = 8.5%). The mean proportion of interpolated pupil data in the analyzed data segments was *M* = 6.0% (*Mdn* = 5.6%, range: 1.0% - 13.7%, *SD* = 3.4%). For the visual control task the mean proportion of the excluded trials was *M* = 2.8% (*Mdn* = 0.0%, range: 0.0% - 42.5%, *SD* = 7.2%). The mean proportion of the interpolated pupil data in the analyzed segments of the used visual trials was *M* = 4.7% (*Mdn* = 4.0%, range: 0.5% - 13.2%, *SD* = 3.0%).

### 2.5. Statistical analysis

As a first step of the statistical analysis, we baseline corrected each individual test trial to quantify the change of the pupil size driven by the cue presentation. The mean pupil size value of the 500 msec period preceding the cue presentation was subtracted from all values of the data segment. All subsequent analyses were done on this baseline corrected data.

As a next step, we used these trial-level baseline corrected data segments to compute participant-level average pupil size curves separately for each participant and each condition. That is, for each participant, we averaged the pupil size curves separately using the trials of the four conditions (*wink, normal, blush* and *hint*).

Then, these participant-level average pupil size curves were used to compare pupillary responses in different conditions for each time point during the presentation of the cue. The comparison was done with Wilcoxon signed-rank tests. To control for multiple comparisons, we used paired-samples cluster based permutation tests, implemented with the help of the MNE-Python software package (Gramfort, 2013; Larson et al., 2023). The approximate z-score of the Wilcoxon test was used as test statistic. The cluster forming threshold was 1.96, which corresponds to the 5% two-sided pointwise significance level. Cluster points were weighted by their test statistics. That is the permutation test considered both the length (duration) of the clusters and the magnitude (z-score) of the difference between the pupil responses at each cluster point.

When assessing the behavioral performance in different experimental conditions we assumed that for a given test trial type the participant gives the correct (or preferred) answer in each test trial independently and with the same probability. Therefore we used binomial tests to compare performances with chance level and Fisher’s exact tests to compare performances in different conditions.

The significance level was 5% (two-sided) for all statistical tests.

## 3. Results

### 3.1. Behavioral task performance

First, we assessed the participants’ behavioral performance in the task, to validate that they correctly understood the instructions and acted accordingly. Figure 2A shows the proportion of correct (rewarded) responses during the test trials for the different conditions.

**Figure 2.**
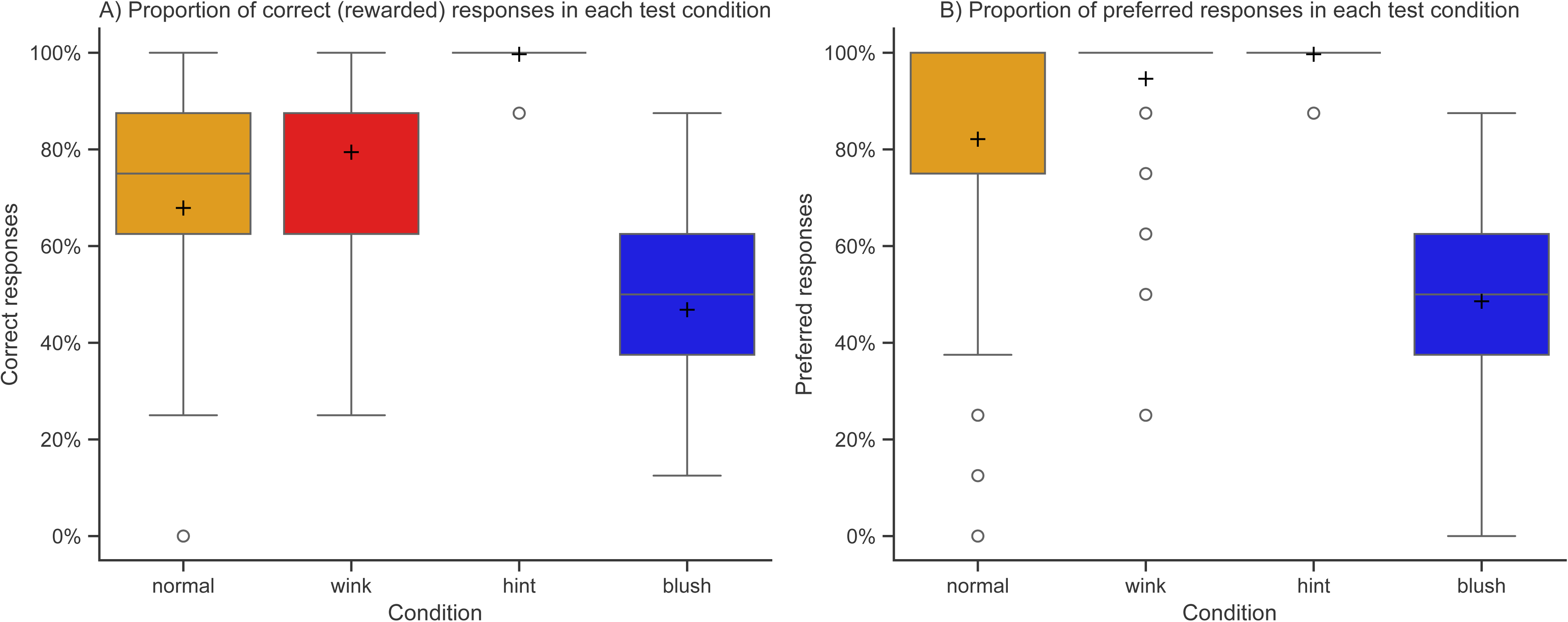
Choices on the test trials. A) Proportion of correct (rewarded) responses during the test trials for each condition B) Proportion of preferred responses during the test trials for each condition. *Note:* +signs indicate mean values.

As expected, the proportion of correct responses was highest in the *hint* condition (*M* = 99.7%, *Mdn* = 100%, range: 87.5% - 100%, *SD* = 2.0%) and lowest in the *blush* condition (*M* = 46.8%, *Mdn* = 50.0%, range: 12.5% - 85.7%, *SD* = 18.5%). The *normal* (*M* = 67.9%, *Mdn* = 75.0%, range: 0.0% - 100%, SD = 23.3%) and *wink* (*M* = 79.4%, *Mdn* = 87.5%, range: 25.0% - 100%, *SD* = 17.5%) conditions had intermediate values. In the *blush* condition participants had no prior knowledge about the rewarded option. In line with this the binomial test showed no significant difference from chance (50%) level for *blush* condition (*p* = .12). In accordance with the instruction and our expectations, in all other conditions, the proportion of correct responses were significantly different (above) chance level (*p* < .001 in all cases).

Fisher’s exact test revealed a significant difference between the proportion of correct responses in the *normal* and *wink* conditions (*p* < .001). It might indicate that some participants did not fully understand the given instructions for one of these conditions. To make sure that the observed differences in pupil responses are not attributable to such possible misunderstanding, we tried to exclude the affected participants. We will show below that excluding participants who tended to select the non-preferred side in either of these conditions and excluding trials where the participant did not select the preferred side does not affect our main results.

Task performance in general, and during *normal* and *wink* trials in particular was highly determined by how well the participants tracked and if they consistently selected the preferred side. Therefore we also calculated the proportions of test trials when the participant selected the preferred direction (Figure 2B). This was also lowest for *blush* trials (*M* = 48.6%, *Mdn* = 50.0%, range: 0.0% - 87.5%, *SD* = 19.4%), in this case the binomial test revealed no significant difference from chance (50%) level (*p* = .50). For *hint* trials the values are identical to the correct response proportions above, as the hinted side always became the (new) preferred option. For *normal* (*M* = 82.1%, *Mdn* = 100%, range: 0.0% - 100%, *SD* = 27.3%) and *wink* (*M* = 94.6%, *Mdn* = 100%, range: 25.0% - 100%, *SD* = 13.1%) trials the values are somewhat lower than for *hin*t trials, but well above chance level (*p* < 0.001 in both cases).

This difference can be attributed to the fact that in the *hint* condition the preferred/rewarded option was explicitly shown to the participant while in *normal* and *wink* conditions it is assumed that the participant has figured out the preferred direction during the filler trials. The Fisher’s exact test also showed a significant difference between the proportions of preferred responses in *normal* and *wink* conditions (*p* < .001). This is in line with the result for the correct response proportions above.

### 3.2. The effect of environmental change on pupil-linked brain arousal

To evaluate the pupil size effect of change we compared the pupillary responses in the *wink* and *normal* test trials (see Figure 3A). As can be seen, the presentation of the cue stimulus evoked an increase in pupil size 1-2 seconds after the cue onset. A large pupil response can be observed also after the disappearance of the cue, indicating that the participants initiated their choice, informed by the cue.

**Figure 3.**
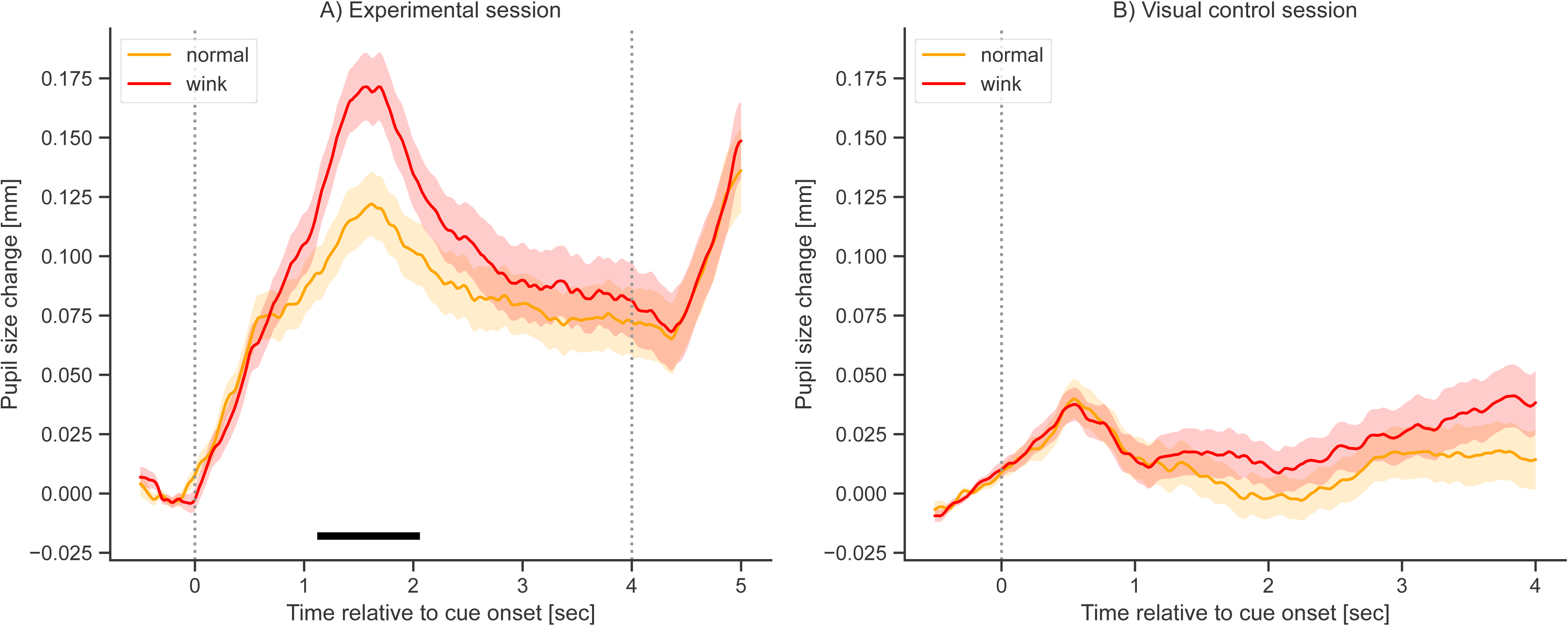
Pupil size changes in the wink and normal conditions. A) Grand-mean pupil dilation curves in mm during cue presentation in the probabilistic reversal task. *Wink* trials are indicated by red, whereas *normal* trials with orange line color. Dashed vertical lines indicate the start of cue presentation and the response period. Horizontal black line indicates significant difference between the conditions (i.e. significant cluster). B) Grand-mean pupil dilation curves in mm during cue presentation in the visual control task. *Wink* trials are indicated by red, whereas *normal* trials with orange line color. The dashed vertical line indicates the start of cue presentation. *Note:* Shading indicates standard error of the mean.

Importantly, for a transient period after stimulus presentation, pupil responses triggered by *wink* trials (see red pupil size curve on Figure 3A) exceed pupil dilations following *normal* trials (see orange pupil size curve on Figure 3A). Wilcoxon signed-rank tests were significant for all time points between 1120-2060 msec, but for no time points after this period. The paired-samples cluster based permutation test showed that this cluster indicated a significant difference between the pupil responses even considering the multiple comparisons (p = .03, see horizontal black line in Figure 3A). Note, that the *wink* and the *normal* conditions differed only in amount of change (see Table 1), thus this difference should reflect the sensitivity of pupil-linked brain arousal to environmental change, and cannot be explained by differences in belief uncertainty.

We then verified that the obtained result is not related to the participants’ observed performance difference between the two conditions (see previous section). To do this we first excluded all subjects from the analysis who did not select the preferred side in more than half of the cases in both *normal* and *wink* test trials. These participants might not properly understood or followed the instructions for one of the trial types. This meant excluding 14 participants (in addition to the one person excluded for poor pupil data), so the remaining sample size was *N* = 65. For the remaining subjects we also excluded all trials where they did not select the preferred direction, which indicated that they deviated from the more advantageous option implied by the task instruction. We then repeated the analysis for the remaining data. The result was very similar to the one before the exclusions. Wilcoxon tests were all significant between 920-2000 msec, but not after this period, with the cluster based permutation test revealing that pupil response was indeed significantly higher for the *wink*, as compared to the *normal* trials (p = .03).

Finally, we also investigated pupil size changes during the visual control task, to evaluate whether there was any difference between the pupil changes evoked solely by the presentation of the different visual cues. The paired-samples cluster based permutation test showed no significant difference between pupil responses evoked by the *wink* and *normal* visual stimuli (Figure 3B). This confirmed that the pupil size effect reported above was indeed related to the performed task and cannot be attributed to differences in the low level visual features of the presented cue stimuli.

### 3.3. The effect of uncertainty on pupil-linked brain arousal

To investigate the effect of uncertainty, we compared pupil response during the *blush* and *hint* test trials. As can be seen on Figure 4A, pupil responses during *blush* trials (see blue pupil size curve on Figure 4A) are larger for most of the trial duration than pupil dilations during *hint* trials (see cyan pupil size curve on Figure 4A). Wilcoxon tests gave significant results for all time points between 1180-5000 msec that is until the end of the analyzed data segments. According to the paired-samples cluster based permutation test this indicated a significant difference between the pupillary responses observed in the *blush* and *hint* conditions even after considering the multiple comparisons (*p* < .001). These conditions had equal amounts of change, but different amounts of uncertainty, thus our results suggest that belief uncertainty exerts a sustained effect on pupil size, and this effect is independent of the detection of environmental change.

**Figure 4.**
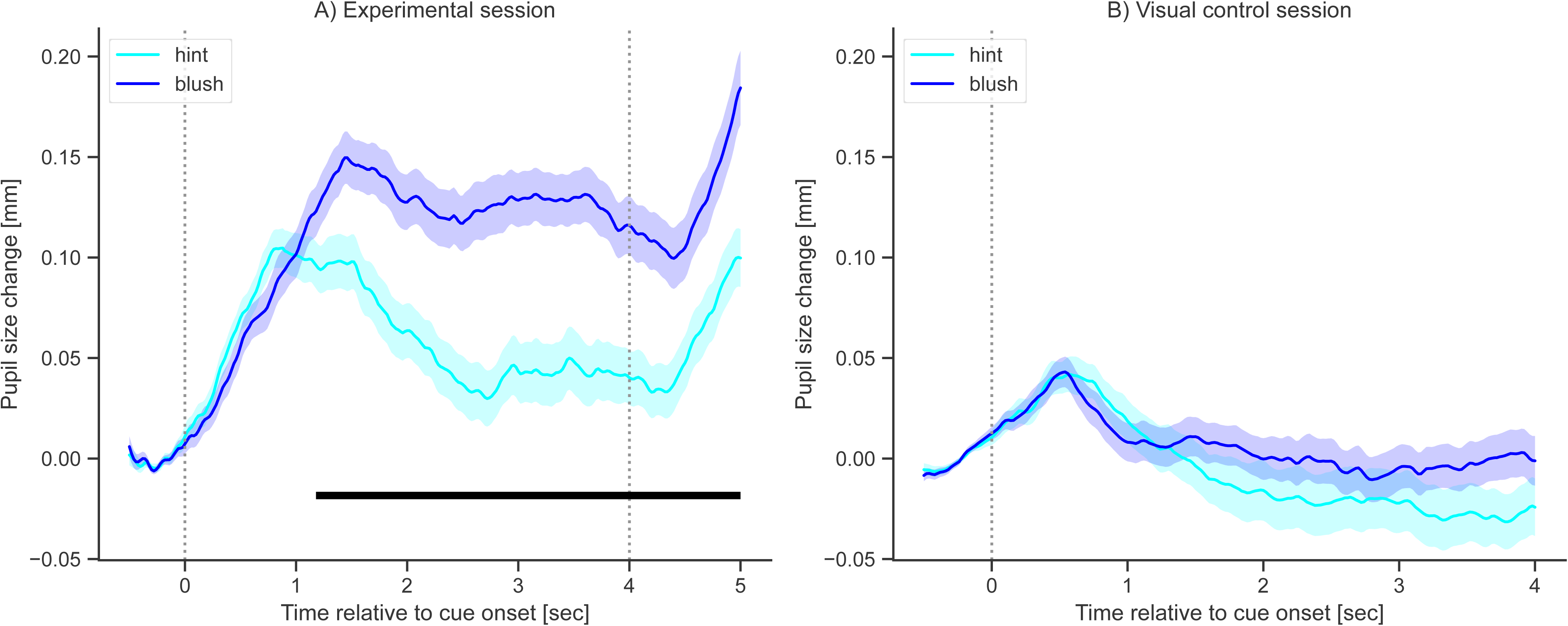
Pupil size changes in the blush and hint conditions. A) Grand-mean pupil dilation curves in mm during cue presentation in the probabilistic reversal task. *Blush* trials are indicated by blue, whereas *hint* trials with cyan color. Dashed vertical lines indicate the start of cue presentation and the response period. Horizontal black line indicates significant difference between the conditions (i.e. significant cluster) B) Grand-mean pupil dilation curves in mm during cue presentation in the visual control task. *Blush* trials are indicated by blue, whereas *hint* trials with cyan color. The dashed vertical line indicates the start of cue presentation. *Note:* Shading indicates standard error of the mean.

Finally we also compared pupil size changes after the presentation of the *blush* and the *hint* cue stimuli during the visual control task (see Figure 4B). No significant difference was revealed by the cluster based permutation test. This suggests that the effect of uncertainty demonstrated during the probabilistic reversal task was not caused by visual differences between the cues.

## 4. Discussion

In our study, we could disentangle the effects of environmental change and belief uncertainty on pupil-linked brain arousal. On the one hand, we demonstrated a short, phasic pupil dilation, which was triggered specifically by a change in environmental regularities, and which was not related to the uncertainty triggered by these changes. On the other hand, we showed that the same magnitude of environmental change triggers different pupil responses, depending on the uncertainty it causes: large uncertainty leads to a more sustained, tonic increase in pupil size.

The effect of environmental change was identified by comparing the *wink* and the *normal* conditions. The posterior probability distributions in these two conditions were characterized by the same entropy (*H* = 0.72), indicating the same amount of uncertainty. However the magnitude of change was large in the *wink* condition (*KLD* = 1.2), whereas no change was present in the *normal* condition (*KLD* = 0). Thus, the larger pupil dilation 1-2 seconds after stimulus onset in the *wink*, as compared to the *normal* condition, reflects the specific effect of change in environmental regularities, because the level of posterior uncertainty was the same in the two conditions. Our result suggests that pupil-linked brain arousal displays a specific sensitivity to the detection of environmental change. This sensitivity manifests itself in a transient phasic response, immediately after the observation of the environmental change. This immediate and transient response profile triggered by the discrepancy between predicted and observed events replicates previous findings showing that pupil size is sensitive to measures of surprise and KL divergence ((Filipowicz et al., 2020; Nassar et al., 2012; Preuschoff et al., 2011; Zénon, 2019). Importantly, the change-related pupil response in our case was not contaminated by the concurrent effects of uncertainty accompanying the surprising information. On a functional level, our results are in-line with theories of the LC/NA system suggesting that pupil-linked brain arousal tracks and signalizes sudden and unexpected changes in the environment, possibly leading to a reconfiguration of cortical networks (Bouret & Sara, 2004; Sales et al., 2019; Yu & Dayan, 2005), and thus this transient dilation can be considered as a signal indicating the need for belief updating.

The specific effect of uncertainty was demonstrated by comparing pupil responses in the *blush* and *hint* conditions. In these two conditions, the amount of change between the prior and the posterior probability distributions was similar (*KLD* = 0.32), but this change resulted in a posterior probability distribution characterized by high (maximal) uncertainty in the *blush* condition (*H* = 1), whereas it led to a probability distribution characterized by low (minimal) uncertainty in the *hint* condition (*H* = 0). This difference in uncertainty caused a significant difference in the evoked pupil response. Pupil dilation was markedly higher when accompanying high (*blush* condition), as compared to low levels of uncertainty (*hint* condition). Importantly, in contrast to the effect of environmental change, this difference between the two conditions was not transient, but sustained: it emerged 1 sec after stimulus onset and remained significant for the remaining time of the trial. Thus, the phasic response following the cue presentation resulted in larger baseline pupil size in the subsequent trial (see the time period of 4-5 sec after cue presentation in Figure 4A). Thus, our results are in line with previous studies showing that higher levels of uncertainty are associated with larger baseline pupil sizes (Filipowicz et al., 2020; Muller et al., 2019; Nassar et al., 2012; Pajkossy et al., 2017). Importantly, in our case, we can be sure that this sustained response is due to uncertainty and is not related to the concurrent effect of surprise or the detection of environmental change. Furthermore, due to the design of our experiment, uncertainty is the result of a single piece of specific information presented to the participants (i.e. the blushed face of the fictional character), and is not a gradually emerging feature of the situation, based on subsequent feedbacks given to previous responses, as was often the case in previous experiments (e.g. Muller et al., 2019; Nassar et al., 2012; Pajkossy et al., 2023). On a functional level, increased belief uncertainty means that the valid environmental regularities have to be identified in the subsequent trial, and so the tonic increase in pupil size might reflect arousal and alertness accompanying this increased cognitive demand of the situation. In line with this, tonic pupil size increase was suggested to accompany exploration in uncertain situations (Hayes & Petrov, 2015; Muller et al., 2019; Pajkossy et al., 2017, 2018) and preparatory increase in baseline pupil size was reported as a sign of task-engagement (Hutchison et al., 2020; Unsworth & Robison, 2016).

On a methodological level, our results emphasize the need for proper experimental control when the psychophysiological, neurobiological or behavioral correlates of belief updating are investigated. With respect to pupil-linked brain arousal, very similar concepts have been linked to its activity (e.g. surprise: (Lavín et al., 2014; Preuschoff et al., 2011; decision uncertainty: De Berker et al., 2016; de Gee et al., 2017; Lavín et al., 2014; Urai et al., 2017; unexpected uncertainty: Pajkossy et al., 2023; belief updating: Filipowicz et al., 2020), with aspects of both change and uncertainty included into the definitions. A good example for this theoretical proximity is that (Yu and Dayan (2005) termed their often cited concept unexpected *uncertainty*, although it measures the probability of a significant environmental change, and thus, in our current framework, it is more related to change than uncertainty. Our results suggest that multiple aspects during the assessment and updating of our beliefs might trigger pupil responses, and when the interrelation of these concepts are not controlled, then one cannot infer what specific aspect of belief updating is driving the observed pupil response (or any other physiological or neural correlate). Note also, that it is usually not sufficient to give a post-hoc estimation for each of these variables on a trial-by-trial basis. In a typical experimental task these concepts strongly correlate and the resulting multicollinearity makes it challenging to disentangle their effects on pupil dilation.

On a theoretical level, distinguishing between these different aspects of the belief updating process might be important, because pupil responses related to environmental change and uncertainty, respectively, might signal the activity of different functional networks in our brain. On the one hand, pupil dilation accompanying the detection of environmental change might reflect the phasic activity of a bottom-up attentional orienting network, detecting significant changes in the environment (Lynn, 1966; Sokolov, 1963; see Corbetta et al., 2008, for a proposal about how the LC/NA system might play a role in this orienting reaction). On the other hand, the more sustained response associated with uncertainty might be related to the activity of a top-down attentional network, which serves to prepare the organism for a period with heightened uncertainty, where new environmental contingencies have to be explored (such role of the LC/NA system was emphasized by Aston-Jones & Cohen, 2005).

In conclusion, we demonstrated that two, often co-occurring decision making variables, environmental change and belief uncertainty, are both associated with pupil-linked brain arousal. The effects are independent in the sense that controlling the value of either of these variables does not eliminate the effect of the other variable. Whereas detecting a change in environmental regularities was associated with a transient pupil dilation, increase in uncertainty was accompanied by a more sustained increase in pupil size. Thus, our results confirm the conclusions of previous reports observing similar dissociations (Filipowicz et al., 2020; Nassar et al., 2012), but due to our experimental design, our conclusions are not influenced by possible correlation between the constructs. Furthermore, our results also raise the possibility that the sensitivity of the pupil responses to environmental change and belief uncertainty, respectively, are linked to the activity of different neural networks.

## Data availability statement

The code for the pupil data preprocessing and the code and data for statistical analyses are publicly available on https://github.com/gesztesigabor/winkorblush.git

